# Pathway-informed Universal Domain Adaptation for Single-cell RNA-seq Data

**DOI:** 10.64898/2026.05.07.723423

**Authors:** Xinrong Wei, Xingyi Li, Huan Liu, Gaoyuan Du, Feng Wei, Xuequn Shang

**Affiliations:** School of Computer Science, Northwestern Polytechnical University, Xi’an,Shaanxi, China; Department of Hepatobiliary and Pancreatic Surgery, General Surgery Center, The First Hospital of Jilin University, Changchun, Jilin, China

**Author notes:** These authors contributed equally to this work (Xinrong Wei, Xingyi Li).

**Keywords:** Single-cell atlases, Universal domain adaptation, Pathway activation, Cell type annotation

## Abstract

The rapid accumulation of single-cell atlases has yielded datasets of unprecedented scale, encompassing samples across diverse platforms, locations and laboratories. This multidimensional complexity drives an urgent need for universal domain adaptation methods capable of achieving precise cell-type annotation. However, existing methods lack computational scalability and fail to integrate biological priors. Here, we develop scPathOT, a pathway-informed universal domain adaptation framework that leverages pathway activation transformations to harmonize single-cell datasets across disparate conditions. We demonstrate the versatility of scPathOT across diverse technological platforms, tissues, disease contexts, cellular senescence and treatment conditions. Crucially, this pathway-informed alignment not only accurately resolves cellular identities but also uncovers functional mechanism. In pancreatic islets, scPathOT delineates a shared stress-repair axis traversed by ***β***–cells prior to their divergence into type 1 and type 2 diabetes-specific states. In aging bone marrow, scPathOT disentangles lineage-specific senescence modules that unexpectedly converge onto unified inflammatory and oxidative-stress programs. Furthermore, application to an in-house pancreatic ductal adenocarcinoma cohort uncovers the mechanistic basis underlying the neoadjuvant chemotherapy-induced reorganization of stromal-immune crosstalk. By coupling biological priors with universal domain adaptation, scPathOT provides a scalable, mechanistically interpretable framework to accelerate biological discovery from atlas-level single-cell data.

## Introduction

Single-cell RNA sequencing now profiles tens of millions of cells across tissues, developmental stages, disease states, and therapeutic contexts, generating large-scale atlases that span heterogeneous biological conditions[1, 2]. Realizing the value of these atlases requires reliable cell-type annotation that remains stable when datasets are integrated across these conditions, since annotation underpins downstream analyses including comparative studies of disease progression, perturbation response, and treatment effect. As single-cell datasets increasingly originate from diverse platforms and encompass strong biological perturbations, achieving consistent annotation across conditions—a problem naturally framed as cross-condition single-cell domain adaptation—has become a persistent task in single-cell data analysis[3].

Cross-condition single-cell domain adaptation poses three challenges that conventional methods struggle to address jointly. First, distribution shifts arise from highly heterogeneous sources: technical variation across sequencing platforms and sample-processing protocols introduces systematic batch effects[4, 5], while disease, aging, and therapy induce condition-specific transcriptional remodeling that reshapes cellular profiles[6]. Methods designed for a single class of shift often do not generalize when distinct shift types are encountered together in atlas-level studies. Second, the cell-type composition between reference and query datasets often differs[7]. Condition-specific subpopulations, transient intermediate states, and rare cell types frequently emerge in the query without counterparts in the reference, while some reference populations are absent in the query; forcing global alignment under this composition mismatch can collapse unrelated cells into the same group—commonly referred to as negative transfer—and distort the underlying biological structure[8]. Third, most existing methods perform integration in raw gene-expression space without incorporating biological prior knowledge. The integration step itself encodes no functional structure, so any biological interpretation depends on bridging from gene-level signals to functional categories through additional, indirect analyses.

Many computational strategies have been developed for single-cell integration and annotation, yet each addresses these challenges only partially. Pathway– or gene set–based representations[9, 10] inject biological prior knowledge by aggregating gene-level signals into functional units, but are not designed to accommodate cross-condition distribution shifts and degrade beyond their training contexts. Mutual nearest neighbor–based methods[11] and variational autoencoders[12, 13] learn general-purpose representations that remove technical batch effects effectively, but typically assume identical cell-type composition between reference and query; under condition-induced transcriptional remodeling they tend to over-correct, collapsing query-specific populations into existing reference clusters. Adversarial domain adaptation approaches[14] introduce flexible nonlinear alignment that handles a broader range of distribution shifts, yet their reliance on global manifold matching obscures biologically meaningful variation when reference and query compositions diverge. Standard supervised annotators[15–27] implicitly assume well-separated cell populations, producing unstable predictions for transitional or condition-specific states; recent ensemble strategies such as popV[28] mitigate individual classifier biases through consensus voting but remain bounded by the closed-set assumptions of their underlying classifiers.

To address these challenges, we develop scPathOT, a pathway-informed universal domain adaptation framework for scRNA-seq data analysis. scPathOT integrates three components, each addressing one of the challenges outlined above. First, each cell is encoded jointly from its highly variable gene expression and from cell–cell similarity graphs constructed on curated pathway-activity spaces, so that biological prior knowledge enters the representation directly rather than only the downstream analyses, providing condition-invariant anchors that ground integration in interpretable functional axes. Second, cross-domain matching is formulated as an unbalanced optimal transport (UOT) problem[8] whose Kullback–Leibler-relaxed marginals do not assume equal cell-type composition between reference and query, allowing query-specific cells to remain unmatched rather than being forced into reference clusters and thereby supporting integration across heterogeneous shift types within a single framework. Third, a lightweight student network learns from UOT-derived soft labels on high-confidence cells and refines predictions on ambiguous boundary cells through entropy minimization, recovering transitional cells along continuous developmental lineages that the conservative UOT alignment would otherwise leave unmatched.

We applied scPathOT to five paradigms that span the heterogeneity encountered in single-cell studies. Two are benchmarking settings: cross-platform integration of peripheral blood mononuclear cell datasets across six sequencing technologies, and large-scale cross-tissue integration of the Tabula Muris atlas covering 20 mouse organs. The remaining three are clinical or physiological scenarios across human and mouse data: pancreatic islets from healthy and diabetic donors, the bone marrow microenvironment across the mouse lifespan, and pancreatic ductal adenocarcinoma biopsies from an in-house cohort comparing patients receiving surgery alone with those receiving surgery combined with neoadjuvant chemotherapy. In these settings, scPathOT produced cross-condition annotations that supported downstream biological analyses at multiple resolutions, including pathway-level dissection of *β*-cell dysfunction in type 1 and type 2 diabetes, gene co-expression network analysis of lineage-specific senescence programs in the aging bone marrow, and characterization of chemotherapy-induced stromal–immune communication remodeling in pancreatic ductal adenocarcinoma.

## Results

### Overview of the scPathOT framework

Integrating single-cell datasets across diverse biological conditions remains a persistent challenge, because conventional methods typically assume identical cell-type composition between reference and query datasets—an assumption that breaks down when distribution shifts and condition-specific cellular populations drive systematic mis-alignment and erroneous annotation. scPathOT addresses this challenge through three components, each targeting one of the issues outlined in the Introduction (Fig. 1a and Supplementary Fig. 1). First, rather than operating on raw expression alone, scPathOT grounds cellular representations in curated biological prior knowledge by constructing multi-view cell–cell graphs whose topology is determined by pathway activation, so that cell–cell proximity reflects functional similarity rather than statistical similarity from gene expression data. Second, cross-condition matching is formulated as an unbalanced optimal transport (UOT) problem, whose Kullback–Leibler-relaxed marginals do not assume equal cell-type composition between reference and query, allowing common cell types to be aligned to source prototypes while query-specific populations remain unmatched rather than being forced into existing reference clusters. Third, a confidence-aware knowledge distillation stage recovers transitional or ambiguous cells that the conservative UOT alignment would otherwise leave unmatched, producing annotation boundaries that respect continuous biological variation along developmental lineages. Together, these components allow scPathOT to operate across the diverse single-cell study designs encountered in practice (Fig. 1b). We evaluate it across five scenarios: cross-platform annotation under platform-induced batch effects, cross-tissue integration of large-scale multi-organ atlases, cross-disease analysis to disentangle shared and disease-specific states, cross-age analysis to trace transcriptional dynamics across lifespan, and cross-treatment analysis to characterize therapy-induced microenvironmental remodeling. Together, these applications position scPathOT as a generalizable framework for cross-condition single-cell data analysis.

**Fig 1.**
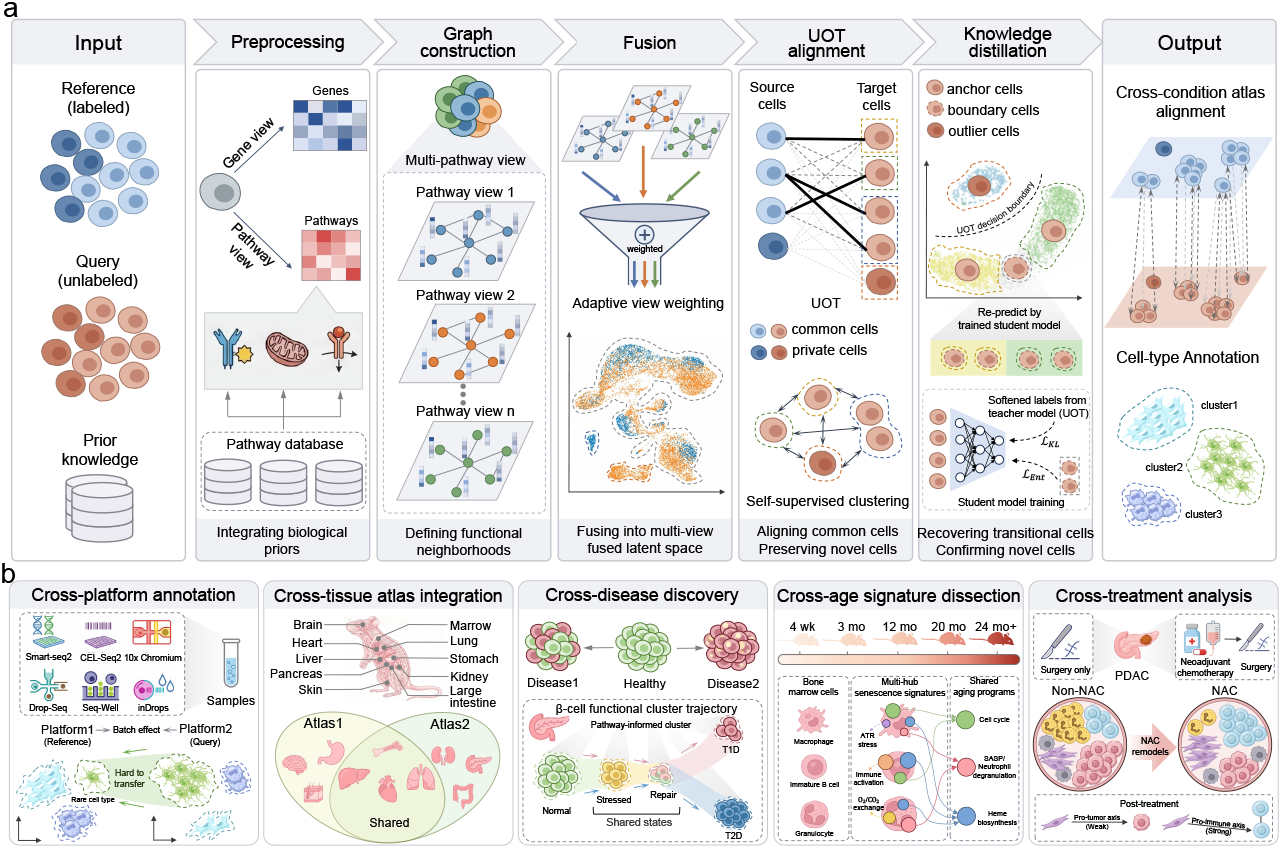
The framework and applications of scPathOT. **a**, scPathOT is a pathway-informed universal domain adaptation framework for cross-condition single-cell integration. Starting from labeled reference and unlabeled query cells, the framework constructs pathway-specific cell-cell graphs to encode functional biological neighborhoods, and fuses multi-pathway representations through a class-conditioned attention module that adaptively weights views per cell, producing a unified latent space. Cross-condition alignment is then formulated as an UOT problem, which aligns common cell types while preserving query-specific populations through marginal relaxation and self-supervised clustering. A subsequent confidence-aware knowledge distillation stage tri-partitions target cells into confident, boundary, and outlier cells, training a student network to recover transitional populations and identify novel cell states. The framework outputs cross-condition atlas alignments and cell-type annotations. **b** scPathOT is evaluated across five biologically distinct scenarios: cross-platform annotation for batch-effect correction across sequencing technologies, cross-tissue atlas integration for harmonizing organ-specific references, cross-disease discovery for isolating shared and disease-specific pathological states, cross-age signature dissection for tracing age-dependent transcriptional dynamics, and cross-treatment analysis for characterizing treatment-induced microenvironmental remodeling.

### Benchmarking scPathOT for Cross-Platform Single-Cell Annotation

To comprehensively evaluate the generalizability and robustness of scPathOT in cross-platform cell-type annotation, we curated 9 peripheral blood mononuclear cell (PBMC) datasets across 6 diverse sequencing platforms[29] (Supplementary Table 1). This compendium enabled the construction of 72 pairwise cross-platform annotation tasks. We benchmarked scPathOT against 21 mainstream single-cell annotation methods (Supplementary Note 1 and Supplementary Table 2). scPathOT consistently outperformed all competing methods across the 72 experimental setups in both average accuracy and macro F1-score (Fig. 2a). Specifically, scPathOT achieved the highest average accuracy of 0.8551, surpassing the second-best method, popV (0.8426), and attained a superior average macro F1-score of 0.7709, followed by Seurat (0.7597). Because accuracy can be inflated by majority-class prediction in long-tailed datasets, we report macro F1 alongside accuracy to weight rare and dominant cell types equally; the macro F1 advantage of scPathOT indicates balanced performance across both dominant and rare cell populations.

**Fig 2.**
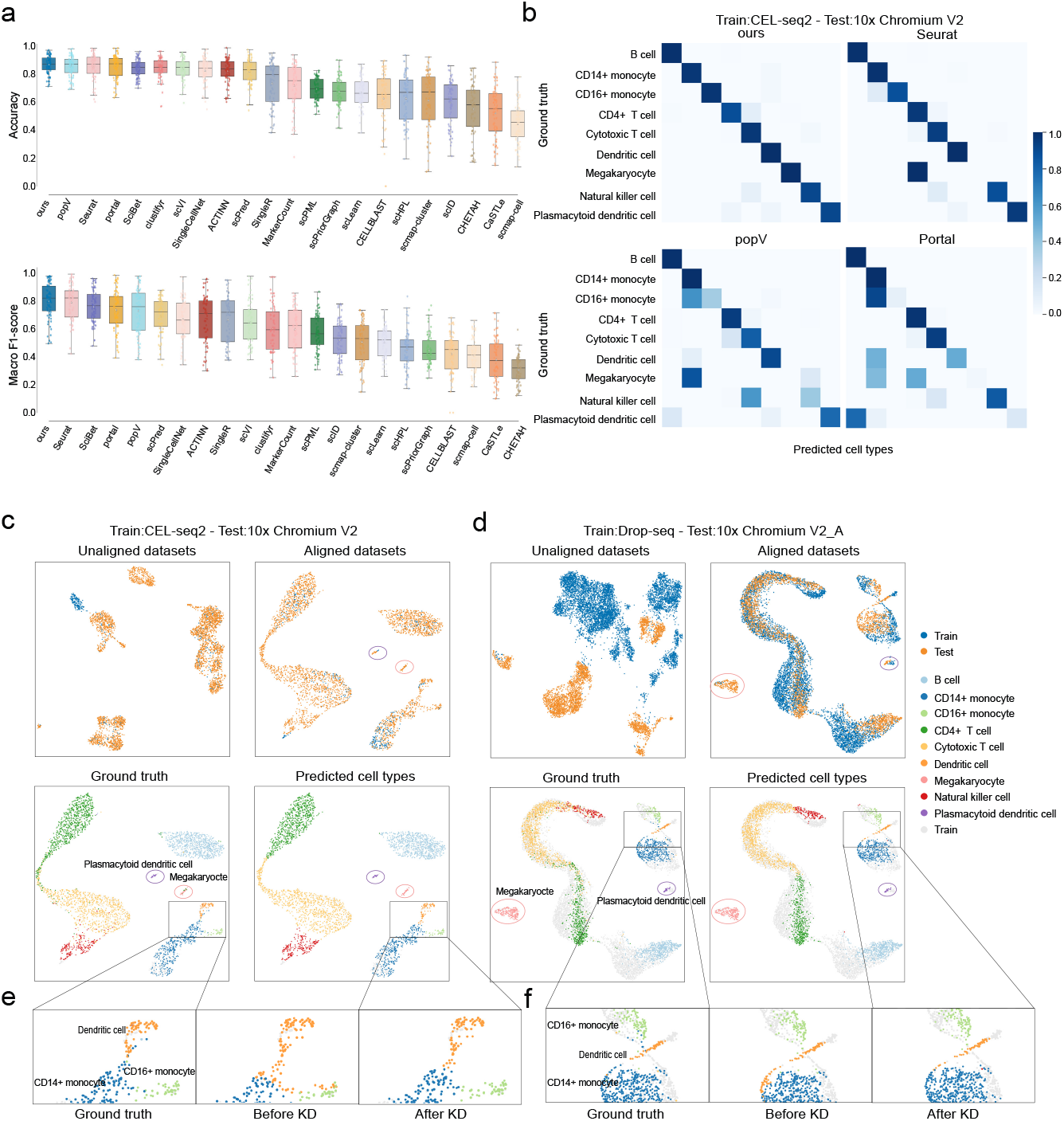
Benchmarking scPathOT for Cross-Platform Single-Cell Annotation. **a**, Boxplots showing the Accuracy (top) and Macro F1-score (bottom) of scPathOT and 21 single-cell annotation tools across 72 cross-platform transfer tasks. **b**, Confusion matrices of cell type predictions by scPathOT, Seurat, popV, and Portal for the transfer task from CEL-seq2 to 10x Chromium V2. **c, d**, UMAP visualizations of the transfer tasks from CEL-seq2 to 10x Chromium V2 (**c**) and from Drop-seq to 10x Chromium V2 A (**d**). For each task, the top panels display the unaligned and aligned datasets, and the bottom panels show the aligned datasets colored by ground truth and predicted cell types. We marked specific regions containing boundary cell types in the bottom panels. **e, f**, UMAP plots of the marked regions in **c** and **d**, respectively. The panels present the ground truth annotations, predictions before knowledge distillation (KD), and predictions after KD.

To further scrutinize the model’s generalization capabilities under extreme data scarcity, we highlight a particularly challenging case from the 72 tasks: predicting 10x Chromium V2 data using a training set composed of merely 463 cells from the CEL-Seq2 platform (Fig. 2b). This case encompasses two rare cell types: plasmacytoid dendritic cells (pDCs) and megakaryocytes. Given that pDCs play a central role in anti-viral immunity and autoimmune diseases[30], and peripheral megakaryocytes (along with their fragments and platelets) are intimately linked to hematological disorders[31], their accurate identification is of profound biological significance. Despite the extreme scarcity of training samples (only one megakaryocyte in the training set), scPathOT achieved accurate feature extraction and transfer. On the test set, scPathOT achieved an accuracy of 0.9490 and a macro F1-score of 0.9412, far exceeding the performance of Seurat (0.8020). The confusion matrix (Fig. 2b) reveals that Seurat erroneously classified all megakaryocytes as CD4+ T cells, entirely failing to capture the defining features of this rare population. In contrast, scPathOT successfully captured these subtle features, enabling precise annotation. Furthermore, UMAP visualization (Fig. 2c) illustrates that scPathOT mitigated cross-platform distribution shifts and aligned the training and test sets within a shared latent space. Notably, the boundaries of rare populations such as pDCs and megakaryocytes remained clearly distinguishable, whereas other baseline methods struggled to maintain this separation (Supplementary Fig. 3). Beyond extreme small-sample scenarios, scPathOT demonstrated equally robust performance on regular-scale datasets. In the experiment using Drop-seq data (6,819 cells) for training and 10x Chromium V2 A data (3,339 cells) for testing, the raw data exhibited massive distribution discrepancies. Following processing with scPathOT, deep integration was achieved between the training and test sets, effectively abolishing cross-platform batch effects(Fig. 2d). Under this setting, scPathOT again ranked first with an accuracy of 0.9383 while preserving rare cell populations.

In single-cell transcriptomic datasets, CD14+ monocytes, CD16+ monocytes, and dendritic cells all arise from the myeloid lineage. They share high transcriptomic similarities and exist along a continuous developmental trajectory[32]. Consequently, their boundaries in the latent space are often blurred, leading to frequent misclassification. To tackle this, we introduced a knowledge distillation (KD) strategy to optimize the initial alignment predictions. Prior to the application of KD, the boundaries of dendritic cells were frequently misassigned to CD16+ monocytes, exacerbating the blurring of classification margins. By leveraging KD, scPathOT uses the soft probability distributions from the teacher to smooth decision boundaries, rendering the predicted distributions closer to the ground truth and effectively correcting misclassifications within continuous developmental lineages(Fig. 2e and Fig. 2f). Quantitative evaluation demonstrated that the integration of the KD module boosted the average accuracy by 0.0199 and yielded a substantial increase of 0.0213 in the average macro F1-score(Supplementary Fig. 4). These results indicate that the KD module improves overall performance and supports finer discrimination among transcriptionally similar cell subtypes.

### Integrating the Large-Scale Cross-Tissue Tabula Muris Atlas

To evaluate scPathOT on large-scale atlases with pronounced compositional imbalances, we integrated the Tabula Muris dataset[33], comprising cells from 20 mouse organs profiled via Smart-seq2 (SS2) and 10x Genomics(Supplementary Note 2). Because the 10x data lacks 7 of the 20 tissues, this presents a “missing cell type” challenge[4] that requires preserving SS2-exclusive populations without forcing erroneous alignments.

Quantitative benchmarking revealed that scPathOT achieved the highest overall performance score (0.661) (Fig. 3a, Supplementary Table 3). scPathOT ranked first in biological-conservation metrics (NMI: 0.886, ARI: 0.832), ahead of the runner-up scVI. Although Portal and popV achieved competitive batch correction, they tended to over-correct, reducing the preservation of biological structure (Supplementary Fig. 5). Conversely, scPathOT effectively eliminated batch effects while maintaining the highest fidelity of cell identities.

**Fig 3.**
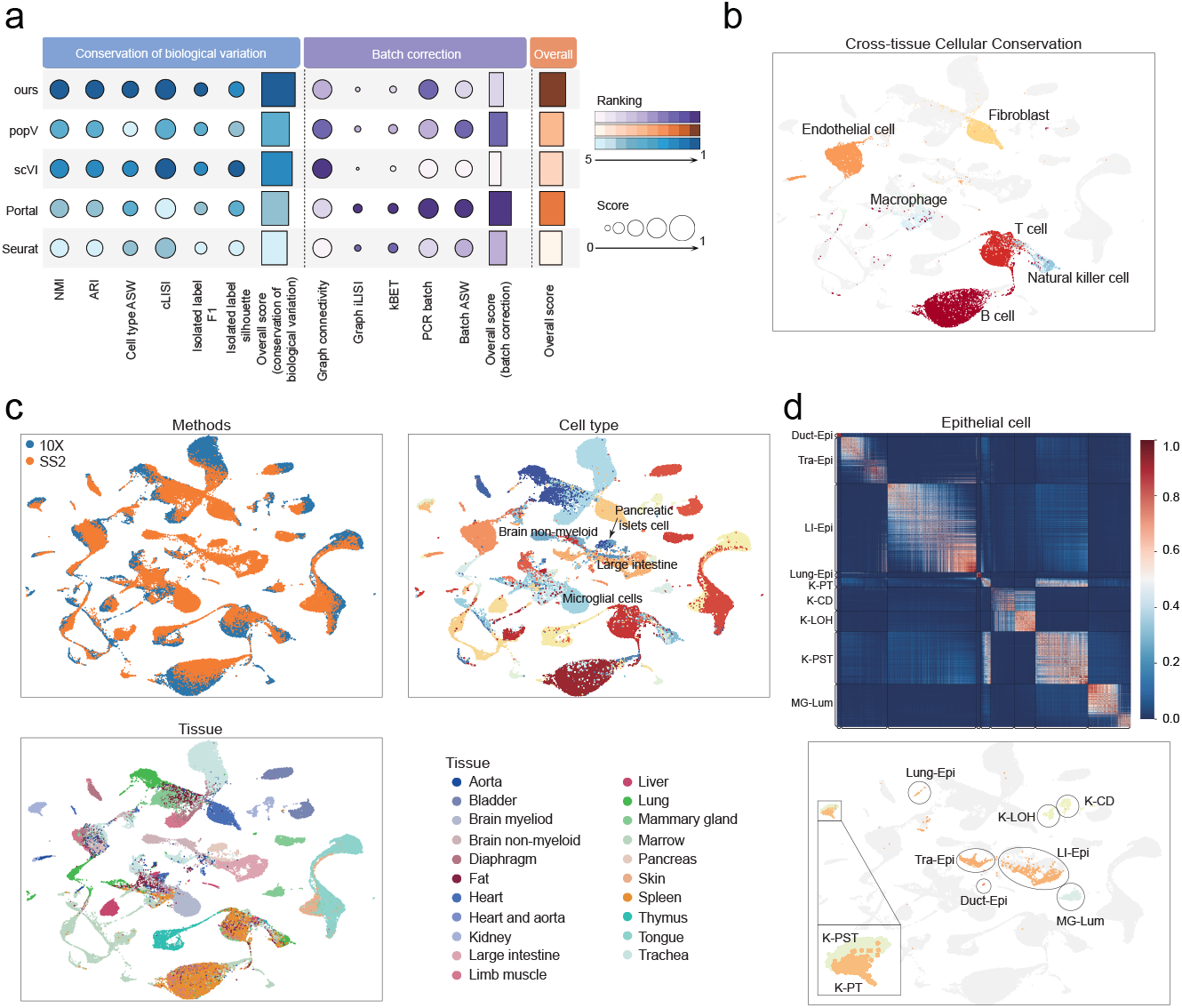
Integrating the Large-Scale Cross-Tissue Tabula Muris Atlas. **a**, Quantitative eval-uation of integration performance. The dot plot compares scPathOT with popV, scVI, Portal, and Seurat across multiple metrics assessing the conservation of biological variation and batch correction. Dot color and size indicate the method ranking and scaled score for each metric, respectively. **b**, UMAP visualization highlighting cross-tissue cellular conservation in the integrated latent space. Major conserved lineages, including immune and stromal cells, are annotated. **c**, UMAP plots of the integrated dataset colored by profiling methods (10x and SS2, top left), cell types (top right), and tissue origins (bottom left). Cell types unique to the SS2 dataset (e.g., brain non-myeloid, pancreatic islets, and large intestine cells) are annotated in the cell type UMAP to illustrate the preservation of dataset-specific populations. **d**, Detailed analysis of epithelial cell subpopulations. The top panel displays a correlation matrix of epithelial cells originating from diverse organs. The bottom panel shows the UMAP projection of these epithelial subpopulations, with zoomed-in views highlighting the structural relationships among kidney-derived epithelial cells. K-PST, kidney proximal straight tubule epithelial cell; K-PT, epithelial cell of proximal tubule; K-LOH, kidney loop of Henle ascending limb; K-CD, kidney collecting duct epithelial cell; Tra-Epi, epithelial cell from trachea; Lung-Epi, epithelial cell of lung; LI-Epi, epithelial cell of large intestine; MG-Lum, luminal epithelial cell of mammary gland; Duct-Epi, duct epithelial cell.

The integrated latent space revealed tight alignment of shared cell populations across tissues and platforms, while tissue-specific cell types formed clearly isolated clusters. For common populations, conserved lineages—including endothelial cells, fibroblasts, macrophages, T cells, natural killer (NK) cells, and B cells—were aligned into tightly cohesive clusters. This integration between 10x and SS2 modalities highlighted the consistent transcriptomic signatures of these immune and stromal cells regardless of their tissue of origin, while preserving their internal biological substructures (Fig. 3b, Fig. 3c left). Importantly, scPathOT achieved this global alignment without erroneously forcing non-overlapping populations into improper clusters. Tissue-specific cell types exclusive to the SS2 dataset, such as brain microglia, pancreatic islet cells, and large intestine epithelia, correctly maintained their unique identities and formed clear, isolated clusters (Fig. 3c right, Supplementary Fig. 6). Collectively, these results demonstrate scPathOT’s ability to disentangle technical batch effects from biological heterogeneity in highly imbalanced datasets.

We further investigated the integration of epithelial cells, a population characterized by its high degree of tissue specificity despite sharing ubiquitous epithelial markers[1]. scPathOT preserved these tissue-specific signatures at high resolution (Fig. 3d). When integrating epithelial cells from diverse organs (kidney, lung, trachea, large intestine, and mammary gland), scPathOT maintained them as mutually independent clusters, structurally congruent with their functional divergence. Notably, kidney proximal tubule (K-PT) and collecting duct (K-CD) cells, which share transcriptomic similarity despite their distinct functions, clustered into adjacent subclusters clearly separated from respiratory or intestinal epithelia.

### Deciphering *β*-Cell Functional Heterogeneity and Disease-Specific States in Diabetes Progression

To evaluate the performance of scPathOT across distinct disease contexts, we integrated scRNA-seq datasets from normal, type 1 diabetes (T1D), and type 2 diabetes (T2D) islets (Supplementary Note 3). By learning exclusively from normal cell features, scPathOT accurately annotated disease-state cells while projecting them into a unified pathway-informed latent space (Fig. 4a). scPathOT achieved the highest annotation accuracy (0.9587) and Macro F1-score (0.8314), exceeding the second-best Portal by 2.06% in accuracy and the second-best popV by 3.3% in macro F1 (Fig. 4b and Supplementary Fig. 7). Given that *β*-cell failure drives both T1D and T2D pathogenesis[34], we focused subsequent analyses on this population. Dimensionality reduction based on the core differentially regulated pathway matrix revealed that normal, T1D, and T2D *β*-cells segregated into distinct functional territories (Fig. 4c), indicating that different disease etiologies induce divergent pathway-level transcriptional programs in *β*-cells[35].

**Fig 4.**
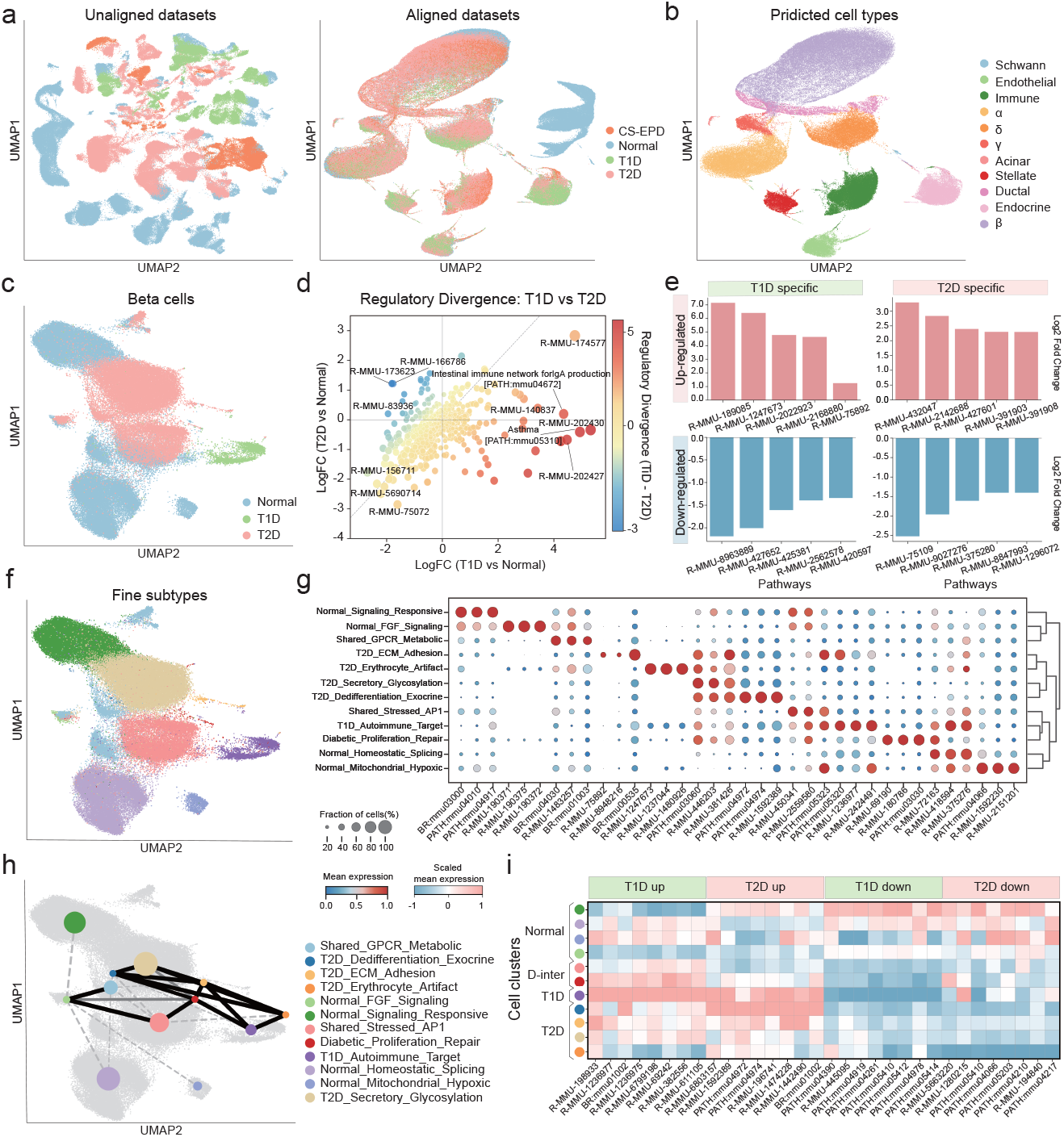
Deciphering *β*-Cell Functional Heterogeneity and Disease-Specific States in Dia-betes Progression. **a, b**, UMAP visualizations of the integrated single-cell transcriptomic datasets comprising Normal, T1D, and T2D models. Panel **a** displays the unaligned and aligned datasets colored by disease conditions, and panel **b** shows the aligned latent space colored by predicted cell types. **c**, UMAP plot of the isolated *β*-cell population, colored by disease conditions. **d**, Scatter plot illus-trating the regulatory divergence between T1D and T2D *β*-cells. The x-axis and y-axis represent the log_2_ fold change (LogFC) of T1D versus Normal and T2D versus Normal, respectively. Point colors indicate the regulatory divergence score (LogFC of T1D minus T2D). **e**, Bar plots showing the LogFC of top up-regulated and down-regulated pathways specific to T1D (left) and T2D (right) *β*-cells. **f**, UMAP plot of fine-grained *β*-cell subtypes identified through high-resolution clustering. **g**, Dot plot showing the expression of top differentially enriched pathways across the fine *β*-cell sub-types. Dot size represents the fraction of cells expressing the pathway, and color intensity indicates the mean expression. **h**, PAGA graph illustrating the connectivity and inferred trajectory among fine *β*-cell subtypes, overlaid on the UMAP embedding. Node colors correspond to the subtypes in **f**, and edge thickness represents connectivity strength. **i**, Heatmap displaying the scaled mean expression of condition-specific pathways (T1D up/down, T2D up/down) across aggregated *β*-cell clusters. T1D, type 1 diabetes; T2D, type 2 diabetes; D-inter, diabetic intermediate; LogFC, log_2_ fold change.

To identify the biological drivers underlying this divergence, we next examined pathways exhibiting shared(Fig. 4d) or disease-specific(Fig. 4e) perturbations. Despite distinct etiologies, both T1D and T2D *β*-cells exhibited a common stress-associated program characterized by increased activity of innate immune pathways, including complement activation (Activation of C3 and C5, R-MMU-174577)[36], together with reduced activity in post-transcriptional regulatory processes such as mRNA editing (R-MMU-75072)[37] and Polo-like kinase–mediated events (R-MMU-156711)[38]. Beyond these common features, disease-specific pathway programs emerged. T1D *β*-cells displayed increased activity in immune synapse–related pathways, including Translocation of ZAP-70 (R-MMU-202430) and Phosphorylation of CD3 and TCR zeta chains (R-MMU-202427). Although these pathways are classically associated with T-cell receptor signaling, their elevated activity in *β*-cells is consistent with enhanced immune engagement within inflamed islets, suggesting that *β*-cells in T1D may engage immune-related signaling cascades within the inflamed islet microenvironment[39]. T1D cells also exhibited increased activity in extracellular matrix remodeling and adhesion-related processes, including DS-GAG biosynthesis (R-MMU-2022923) and Nectin/Necl trans heterodimerization (R-MMU-420597), consistent with structural disruption during insulitis[40]. In contrast, T2D *β*-cells showed increased activity in lipid inflammatory mediator pathways, including Synthesis of 5-eicosatetraenoic acids (R-MMU-2142688) and Prostanoid ligand receptors (R-MMU-391908), both associated with *β*-cell dysfunction under metabolic stress[41]. T2D cells additionally displayed altered osmotic and ion transport regulation, reflected by increased activity of Aquaporins (R-MMU-432047) and SLC26 transporters (R-MMU-427601), suggesting perturbed cellular homeostasis[42]. Consistent with impaired *β*-cell function, these cells also exhibited reduced activity in anabolic programs such as Triglyceride biosynthesis (R-MMU-75109) and altered regulation of voltage-gated potassium channels (R-MMU-1296072), which may disrupt membrane excitability and glucose-stimulated insulin secretion[43].

To further dissect *β*-cell heterogeneity, we performed high-resolution clustering based on pathway activity profiles (Supplementary Note 4), identifying specialized functional subpopulations (Fig. 4f). By evaluating the top differentially activated pathways, we assigned clear biological identities to these fine subtypes (Fig. 4g and Supplementary Note 5). Normal *β*-cells were dominated by homeostatic subsets such as Normal Signaling Responsive, characterized by high activity in MAPK signaling (PATH:mmu04010) and Prolactin signaling (PATH:mmu04917), pathways important for *β*-cell survival and plasticity. T1D samples were predominantly composed of a T1D Autoimmune Target population displaying increased activity in Autoimmune thyroid disease (PATH:mmu05320) and DAP12 signaling (R-MMU-2424491), consistent with inflammatory stress. T2D samples contained a distinct T2D Dedifferentiation Exocrine subtype marked by increased activity in Pancreatic secretion (PATH:mmu04972) and Protein digestion and absorption (PATH:mmu04974), indicating loss of mature endocrine features and acquisition of alternative functional programs[44, 45]. Additional T2D-specific populations included T2D Secretory Glycosylation, associated with increased protein export activity (PATH:mmu03060), consistent with elevated secretory stress, and T2D Erythrocyte Crosstalk, characterized by activation of erythrocyte-associated gas exchange pathways (R-MMU-1247673). This transcriptional signature suggests ectopic activation of erythrocyte-associated transcriptional programs in stressed *β*-cells, the functional significance of which warrants further investigation[46].

To connect these functional subtypes, we reconstructed a pathway-informed trajectory using PAGA (Fig. 4h). While PAGA infers topological rather than temporal relationships, this topology suggests that homeostatic and terminal disease states were linked through a shared intermediary axis rather than direct transitions. This axis was anchored by a Shared Stressed AP1 population exhibiting elevated activity of AP-1 transcription factor activation (R-MMU-450341) and oxidative stress–induced senescence pathways (R-MMU-2559580)[47], suggesting that *β*-cells from different disease contexts may converge onto a common stress-responsive program. Cells within this state subsequently transitioned into a shared Diabetic Proliferation Repair node characterized by increased activity in DNA replication (PATH:mmu03030) and telomere extension pathways (R-MMU-69190), consistent with a transient compensatory response. Following this intermediate phase, the trajectory bifurcated into disease-specific branches (Fig. 4i and Supplementary Note 6). Along the T1D branch, pathway activity progressively increased for antigen presentation (R-MMU-1236975) and immunoregulation interactions (R-MMU-198933), accompanied by reduced activity of cell adhesion molecules (PATH:mmu04514), consistent with enhanced immune interaction during autoimmune-mediated *β*-cell loss[48]. Conversely, the T2D branch showed progressive activation of exocrine-associated pathways (PATH:mmu04972, PATH:mmu04974) and extracellular matrix remodeling (R-MMU-1592389), together with reduced activity of apoptosis (PATH:mmu04210) and necroptosis (PATH:mmu04217). These dynamics indicate that *β*-cells in T2D acquire alternative transcriptional states rather than activating canonical cell death programs under chronic metabolic stress[49]. Together, these analyses delineate a unified progression model of *β*-cell dysfunction. *β*-cells from both disease contexts first converge on a shared stress-associated state and transiently engage compensatory repair programs, before diverging into immune-interacting states in T1D versus identity-remodeled adaptive states in T2D.

### Characterizing Lineage-Specific and Convergent Senescence Programs in the Aging Bone Marrow Microenvironment

In vivo aging introduces high transcriptional noise and cellular heterogeneity, both of which complicate cross-age data integration. We applied scPathOT to a mouse bone marrow atlas spanning 4 weeks to over 20 months and benchmarked it against popV, Portal, scVI, and Seurat. scPathOT achieved the highest Accuracy and Macro F1-score across the four baselines (Fig. 5a, Supplementary Fig. 8). UMAP projections confirmed that scPathOT mitigated cross-condition shifts associated with development and aging, aligning datasets across the 4-week to 20-month lifespan (Fig. 5b), while effectively preserving the cohesiveness and biological topology of the predicted cell type clusters (Fig. 5c). Notably, mouse bone marrow data encapsulates a highly complex hematopoietic hierarchy, where cellular differentiation is a highly continuous process (from hematopoietic stem cells to various progenitors and mature cells), characterized by numerous elusive intermediate transition states[50]. Furthermore, the aging process is accompanied by a pronounced myeloid bias[51], which further distorts the cellular manifold. In this challenging continuous manifold alignment task, Portal, which depends heavily on manifold alignment, showed lower performance under this regime (Macro F1 = 0.6048). This is likely because forcing alignment across distant age batches along continuous differentiation trajectories can over-correct and obscure transitional cell states. The ensemble-based popV also achieved lower performance (Macro F1 = 0.7791). These results indicate that scPathOT captures age-invariant latent representations while preserving continuous differentiation trajectories.

**Fig 5.**
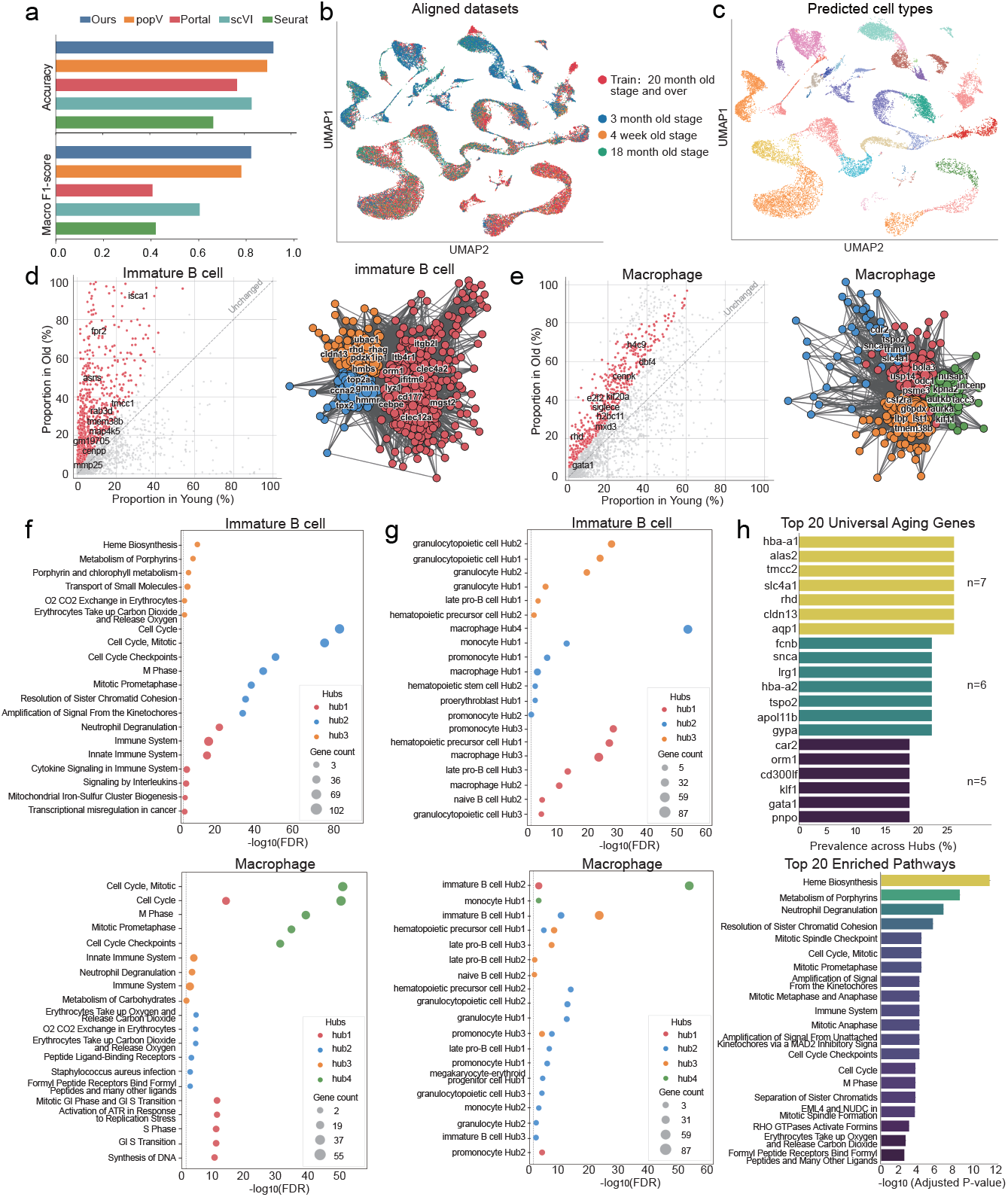
Characterizing Lineage-Specific and Convergent Senescence Programs in the Aging Bone Marrow Microenvironment. **a**, Bar plots comparing the Accuracy (top) and Macro F1-score (bottom) of scPathOT with popV, Portal, scVI, and Seurat for the integration of the aging bone marrow dataset. **b, c**, UMAP visualizations of the integrated dataset, colored by age stage (**b**) and predicted cell types (**c**). **d, e**, Identification of age-dynamic genes and their co-expression networks in immature B cells (**d**) and macrophages (**e**). In each panel, the left scatter plot compares the proportion of cells expressing specific genes in young versus old mice, with red dots highlighting genes with significant age-dependent expression changes. The right network graph illustrates the co-expression relationships among these genes, where nodes represent individual genes and colors denote distinct gene modules (Hubs). **f**, Dot plots showing the functional enrichment of pathways for the identified Hubs in immature B cells (top) and macrophages (bottom). The x-axis represents the − log_10_ (FDR). Dot size indicates the number of genes, and color corresponds to the specific Hubs. **g**, Dot plots illustrating cross-lineage hub associations for immature B cells (top) and macrophages (bottom). The y-axis lists gene modules from various cell types, showing their enrichment significance against the specific Hubs. **h**, Bar plots displaying the top 20 universal aging genes ranked by their prevalence across all cell-type-specific hubs (top), and the top 20 universally enriched pathways ranked by − log_10_ (Adjusted P-value) (bottom).

Cellular senescence is heterogeneous and typically arises in only a minor fraction of cells within aged tissues[52]. Consequently, traditional differential expression analyses are often confounded by global organismal aging or shifts in population-specific proliferation states. Inspired by the analytical framework of SenePy[53], we evaluated age-dynamic genes based on the proportional increase of expressing cells rather than relying solely on fold-change (Supplementary Fig. 9). By constructing co-expression networks of age-dynamic genes, we observed that the senescence signature of a single cell type is composed of multiple topologically segregated gene centers (Hubs) (Fig. 5d,e). Among the 24 mouse bone marrow cell types, 15 contained multiple gene subnetwork clusters, yielding a total of 32 independent senescence signatures (Supplementary Fig. 10). Functional enrichment analysis of these hubs (Fig. 5f, Supplementary Fig. 11) revealed that distinct gene hubs within individual cell types orchestrate divergent senescence programs. For instance, bone marrow macrophages exhibited four functionally distinct senescence hubs. Hub 4 was characterized by “Mitotic Cell Cycle” pathways, consistent with alterations in proliferative capacity. In contrast, Hub 1 highlighted an abortive or dysregulated cell cycle state associated with DNA damage[54], suggested by enrichment for “G1/S Transition” and “Activation of ATR in Response to Replication Stress”. Beyond cell cycle dysregulation, Hub 3 captured the Senescence-Associated Secretory Phenotype (SASP), dominated by “Innate Immune System” and “Neutrophil Degranulation” signatures. Hub 2 revealed a non-canonical stress-response program; its association with “O2/CO2 Exchange in Erythrocytes” and “Formyl Peptide Receptors” likely reflects the compensatory upregulation of antioxidant genes like hemoglobin to combat niche oxidative stress, alongside the sensing of mitochondria-derived Damage-Associated Molecular Patterns (DAMPs) via formyl peptide receptors[55]. This modularity was similarly observed in immature B cells. Here, distinct hubs independently governed immune activation (Hub 1) and cell cycle arrest (Hub 2), while a third module (Hub 3) was dedicated to “Heme Biosynthesis”, a critical antioxidant metabolic reprogramming mechanism[56]. Together, these results indicate that bone marrow cell aging involves multiple parallel processes governed by functionally distinct gene modules—encompassing cell cycle dysregulation, SASP-associated signaling, replication stress responses, and metabolic adaptation—rather than a single uniform aging program.[57].

Although senescence triggers vary across different hematopoietic lineages in the bone marrow, we next investigated whether these programs converge onto shared stress-response and inflammatory pathways during late-stage aging. Cross-lineage hub association analysis revealed extensive co-enrichment of transcriptional programs across functional hubs of different cell types (Fig. 5g, Supplementary Fig. 12). Among these, Immature B cell Hub 1 exhibited the highest degree of cross-lineage overlap, showing significant gene sharing with Macrophage Hub 3, Promonocyte Hub 3, and Hematopoietic Precursor Cell Hub 1 (− log_10_(FDR) *>* 20 for all three pairs). Macrophage Hub 3 further overlapped with Hematopoietic Precursor Cell Hub 1 and Late Pro-B cell Hub 3 (− log_10_(FDR) *>* 10). Given that Immature B cell Hub 1 and Macrophage Hub 3 are both associated with inflammatory immune activation and SASP-related signaling, this convergence suggests that lymphoid and myeloid lineages share partially overlapping inflammatory secretion programs during aging[58, 59], despite their distinct developmental origins. To further explore this commonality, we extracted genes from all bone marrow cell-type signatures. While no single gene was present in every signature, many genes were statistically overrepresented across cross-lineage signatures. Among recurrent genes, hemoglobin subunit genes (e.g., *Hba-a1*), canonically expressed in erythroid cells, appeared in the age-dynamic gene sets of up to 7 non-erythroid cell types, suggesting ectopic antioxidant responses as a shared feature of bone marrow aging[60]. Carbonic anhydrase genes (e.g., *Car2*) were similarly recurrent across 5 cell types, consistent with dysregulation of intracellular pH homeostasis under aging-associated stress[61]. Pathway enrichment of these high-frequency genes (Fig. 5h) revealed that “Heme Biosynthesis”, “Neutrophil Degranulation” and “Cell Cycle Checkpoints” ranked among the top shared pathways. These pathways are consistent with heme-dependent oxidative stress adaptation, SASP-driven degranulation-associated inflammatory signaling[62], and age-related decline in stem/progenitor proliferative capacity via cell cycle checkpoint activation[63]—though precise mechanistic contributions warrant further functional validation. Taken together, our analyses reveal that although bone marrow cells undergo aging through heterogeneous lineage-specific programs, these programs share common pathway-level signatures centered on oxidative stress adaptation, inflammatory secretion, and cell cycle dysregulation—a pattern consistent with the partial cross-cell-type pathway commonality observed in broad multi-tissue single-cell analyses[53].

### Unraveling Neoadjuvant Chemotherapy–Associated Remodeling of the Tumor Immune Microenvironment in Clinical PDAC

To systematically evaluate the impact of neoadjuvant chemotherapy (NAC) on the tumor microenvironment (TME), we applied scPathOT to integrate and annotate single-cell RNA sequencing data from a “surgery-only” patient (Non-NAC) and a “surgery + NAC” patient (NAC). Dimensionality reduction and visualization demonstrated that scPathOT effectively mitigated inter-sample batch effects, achieving coherent alignment of the two samples within the manifold space (Fig. 6a), with clearly delineated boundaries for each cell subpopulation (Fig. 6b). Cell-type annotations derived from canonical marker genes in the Non-NAC sample exhibited highly consistent expression patterns in the NAC sample (Fig. 6c), confirming the robustness of the classification approach. Quantitative analysis of cellular proportions revealed structural remodeling of the TME following NAC (Fig. 6d, Supplementary Table 4): compared to the Non-NAC sample, the proportion of neutrophils declined markedly (from 29.3% to 3.5%), whereas T cells (from 15.5% to 34.0%) and fibroblasts (from 14.3% to 21.6%) expanded. These compositional shifts reflect a macroscopic reorganization of the TME, suggesting that surviving resident cells likely underwent transcriptional adaptations in response to chemotherapeutic stress—an observation that motivates the cell-type-specific molecular analyses below.

**Fig 6.**
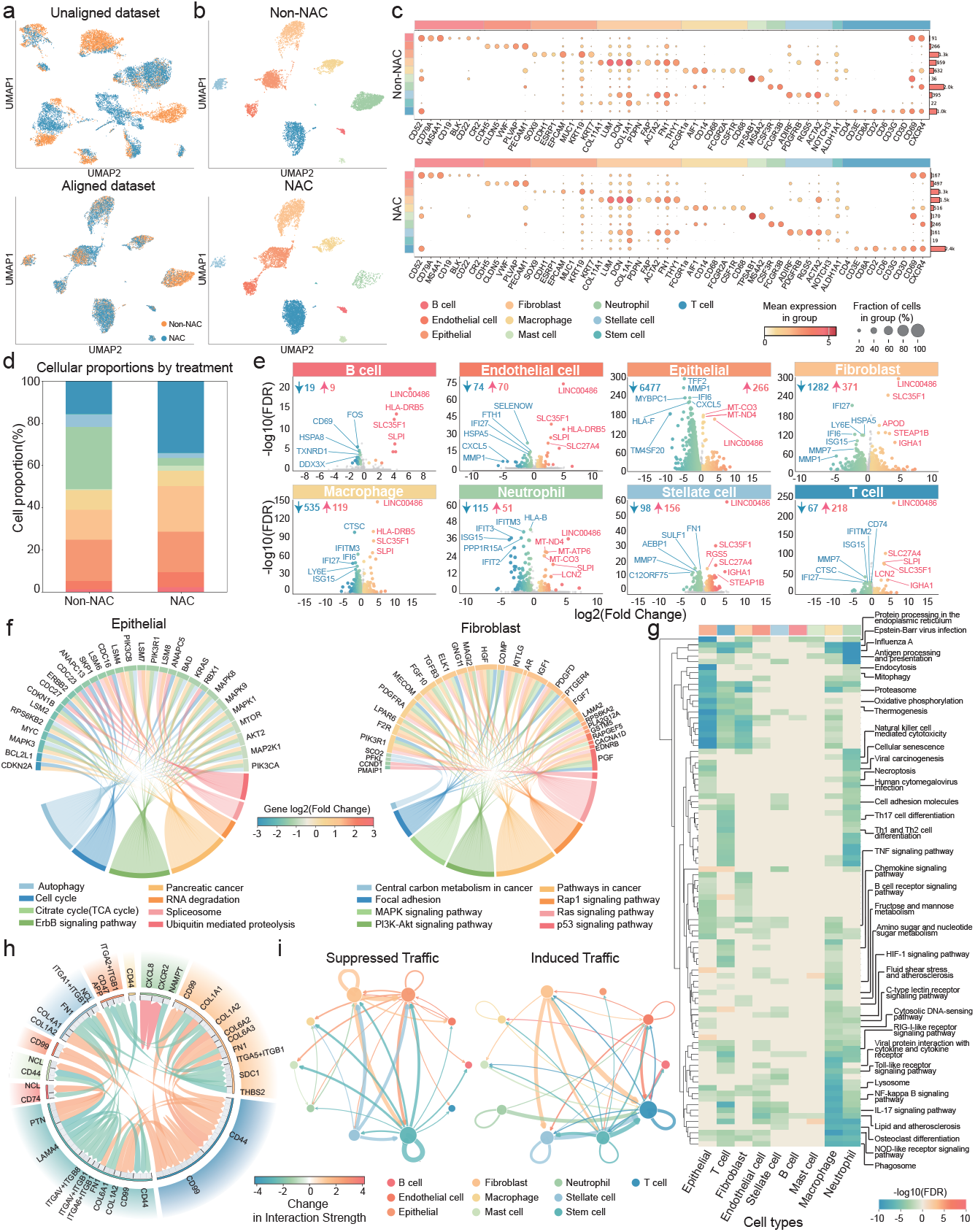
Unraveling Neoadjuvant Chemotherapy–Associated Remodeling of the Tumor Immune Microenvironment in Clinical PDAC. **a, b**, UMAP visualizations of the integrated single-cell transcriptomic datasets from Non-NAC and NAC samples. **a**, The unaligned (top) and aligned (bottom) datasets colored by treatment condition. **b**, Aligned cells separated by Non-NAC (top) and NAC (bottom) conditions, colored by annotated cell types. **c**, Dot plot showing the expression of canonical marker genes across identified cell types in Non-NAC (top) and NAC (bottom) groups. Dot size represents the fraction of expressing cells, and color intensity indicates the mean expression level. **d**, Stacked bar chart comparing the cellular proportions of each cell type between the Non-NAC and NAC samples. **e**, Volcano plots illustrating differentially expressed genes (DEGs) between NAC and Non-NAC conditions across eight major cell types. The x-axis represents log_2_(fold change), and the y-axis represents − log_10_(adjusted P-value). **f**, Chord diagrams displaying the enrichment of selected cell-type-specific functional pathways for epithelial cells (left) and fibroblasts (right). The upper arcs represent the top 30 DEGs (colored by regulation direction), and the lower arcs represent a subset of enriched pathways unique to each cell type. Arc lengths indicate the number of involved pathways per gene or genes per pathway, with inner chords connecting genes to their respective pathways. **g**, Heatmap showing the directional enrichment of shared pathways across different cell types. Colors indicate the regulation direction and significance. **h**, Chord diagram detailing the top 40 most altered ligand–receptor interactions. The outer colored sectors represent cell types, with arc lengths proportional to their total interaction involvement. The inner grey bands denote specific ligands or receptors. Chord colors indicate the change in interaction strength. **i**, Network graphs illustrating the top 100 suppressed (left) and top 100 induced (right) cell–cell communication events following NAC treatment. Nodes represent cell types, with sizes proportional to their altered communication activity. Edge thickness indicates the magnitude of change in interaction strength.

At the transcriptional level, differentially expressed gene (DEG) analysis (Fig. 6e, Supplementary Table 5) revealed extensive cellular reprogramming across the TME induced by NAC. Broadly, all major cell types exhibited extensive transcriptional changes following NAC, with downregulated genes predominating across multiple immune and stromal subpopulations. Among the consistently upregulated transcripts across cell types, the long non-coding RNA *LINC00486* and the solute carrier family member *SLC35F1* were most prominently shared. Given that *LINC00486* has been reported to function as a tumor-suppressive lncRNA that engages NKRF– NF-*κ*B/TNF-*α* signaling[64] and has recently been implicated in injury-responsive transcriptional regulation, while *SLC35F1* is a solute transporter whose deficiency has been linked to cellular degeneration and impaired tissue function in aging models[65], their concordant pan-cellular upregulation suggests a potential stress-associated adaptive response in surviving TME cells under chemotherapeutic pressure. Nevertheless, whether these genes directly mediate chemotherapy tolerance or merely mark stress-exposed cell states will require functional validation through gene-knockout or perturbation studies. Additionally, the downregulation of matrix metalloproteinases (*MMP1, MMP7*) and the chemokine *CXCL5* in epithelial cells and fibroblasts is consistent with attenuation of extracellular matrix (ECM) remodeling and reduced myeloid recruitment capacity[66]. Concurrently, the downregulation of interferon-stimulated genes (ISGs; e.g., *ISG15, IFI27*) in T cells and macrophages likely reflects broad suppression of interferon signaling following chemotherapy[67]; whether this represents resolution of chemotherapy-induced interferon signaling or broader suppression of immune activation warrants further investigation.

Pathway enrichment analysis linked these transcriptional changes to specific functional programs. Cell-type-specific network analysis (Fig. 6f, Supplementary Fig. 13) revealed that NAC was associated with divergent transcriptional responses across distinct subpopulations. In epithelial (tumor) cells, core oncogenic drivers[68, 69]— including *ERBB2, KRAS, MYC*, and PI3K–AKT–mTOR pathway components (*PIK3CA, AKT2, MTOR*)—were significantly downregulated, with concurrent suppression of proliferation-related gene sets such as “Cell cycle” and “ErbB signaling pathway”, consistent with direct cytotoxic and anti-proliferative effects of NAC on tumor cells[70]. In contrast, fibroblasts exhibited a distinct transcriptional profile following NAC, characterized by upregulation of multiple growth factor and receptor genes (e.g., *PDGFRA, TGFB3, HGF, FGF7* /*FGF10*) alongside transcriptional signatures associated with “Focal adhesion”, “MAPK signaling” and “PI3K–Akt signaling” pathways. This pattern is consistent with a chemotherapy-induced stromal wound-healing response to cytotoxic tissue damage[71], and may underlie the subsequent redirection of collagen- and fibronectin-mediated signaling observed in the cell–cell communication analysis, away from pro-tumor (fibroblast→tumor, via *ITGA2* /*ITGB1*) and toward pro-immune (fibroblast→T cell, via *CD44*) axes.

Beyond cell-type-specific responses, NAC induced a convergent and broadly suppressive transcriptional effect across the TME (Fig. 6g, Supplementary Table 6). Profiling of pathways shared across multiple cell types revealed that the majority of shared pathways showed consistent downregulation across multiple cell types. This widespread transcriptional suppression was concentrated in three core biological modules. First, broad attenuation of inflammatory and innate immune signaling was observed; innate immune pathways (NOD-like, RIG-I-like, and Toll-like receptor signaling), pro-inflammatory cascades (NF-*κ*B, IL-17, and TNF signaling), and “Antigen processing and presentation” were downregulated across 5–7 distinct cell types, including epithelial cells, fibroblasts, macrophages, and T cells. These changes are consistent with resolution of chemotherapy-induced cytotoxic inflammation across the residual TME, rather than reflecting broad immune suppression. Second, a global reduction in metabolic activity was evidenced by the consistent attenuation of “Oxidative phosphorylation”, “Fructose and mannose metabolism”, “Glycolysis / Gluconeogenesis”, and “Lipid and atherosclerosis” gene sets. Third, suppression of “Protein processing in the endoplasmic reticulum”, “Proteasome”, “Lysosome”, and “Phagosome” pathways across multiple cell types indicated a coordinated attenuation of proteostatic machinery, consistent with reduced intracellular protein turnover. Collectively, this pan-cellular transcriptional suppression suggests that chemotherapy drives the TME toward a broadly quiescent state characterized by diminished metabolic activity, reduced inflammatory signaling, and attenuated proteostatic machinery—a pattern that may reflect an adaptive response by resident TME cells to severe cytotoxic stress, setting the stage for the stromal–immune communication remodeling described below.

Finally, intercellular communication analysis revealed that NAC reshaped the signaling architecture of the TME. NAC was associated with attenuation of several pro-tumorigenic communication axes involving fibroblasts and epithelial cells (Fig. 6h, Supplementary Table 7). For instance, *COL1A1* /*COL1A2* –(*ITGA2* +*ITGB1*) collagen–integrin signals originating from fibroblasts were weakened, consistent with reduced ECM-mediated support for tumor cells. Simultaneously, the *THBS2* –*CD47* interaction—a pro-tumorigenic axis that activates MAPK/ERK5 signaling in tumor cells[72]—was suppressed, consistent with disruption of stromal–tumor cooperation in immune escape. In contrast, NAC was associated with the emergence of a T cell-centered intercellular communication network (Fig. 6i). Fibroblasts showed increased output of *COL1A1* /*FN1* –*CD44* signals, consistent with enhanced capacity to support T cell adhesion and tissue infiltration[73]. Additionally, *CD99* –*CD99* homophilic interactions were elevated among T cells, endothelial cells, and fibroblasts, a pattern associated with facilitation of leukocyte transendothelial migration[74]. Taken together, these findings suggest that NAC not only suppresses tumor cell proliferation but is also associated with ECM remodeling, broad dampening of microenvironmental inflammation, and reorganization of intercellular communication networks in ways potentially favorable to immune surveillance—observations that may provide a mechanistic rationale for exploring NAC-based combination strategies with immunotherapy[75], though these single-sample observations require replication in larger cohorts.

## Discussion

Single-cell atlases now span platforms, tissues, diseases, lifespans, and treatments, but existing domain adaptation methods for single-cell data are typically developed for one type of shift at a time, treat cells as featureless points, and leave biological prior knowledge unexploited, limiting both their generalizability across heterogeneous conditions and the interpretability of the integrated representation. Here we introduced scPathOT, a pathway-informed universal domain adaptation framework that grounds cell representations in curated pathway-activity spaces, formulates cross-domain matching as an unbalanced optimal transport problem with Kullback– Leibler-relaxed marginals to accommodate cell-type composition mismatch between reference and query, and recovers transitional cells through a confidence-aware knowledge distillation stage. By coupling biological priors with universal domain adaptation in a single framework, scPathOT supports cross-platform, cross-tissue, cross-disease, cross-age, and cross-treatment analysis. Beyond cell-type annotation, it enables high-resolution biological discovery—charting a shared stress–repair axis along which T1D and T2D *β*-cells converge before diverging into disease-specific terminal states, disentangling lineage-specific senescence modules that converge onto common inflammatory and oxidative-stress signatures in the aging bone marrow, and resolving the reorganization of stromal–immune communication in chemotherapy-treated pancreatic ductal adenocarcinoma.

Despite these strengths, scPathOT has several limitations. First, although scPathOT can discover query-specific cell populations without supervision, source-side annotation is still required to anchor common-class alignment; performance may therefore degrade in biological contexts lacking high-quality reference cell-type labels, such as understudied tissues or rare disease settings. Second, the pathway-informed graph construction depends on curated pathway databases (e.g., KEGG, Reactome), which may provide limited coverage for non-model organisms or emerging disease contexts; incorporating data-driven gene co-expression modules could mitigate this dependency in future extensions. Third, scPathOT’s behavior under extreme domain shifts—particularly cross-species integration, where orthologous genes may exhibit divergent expression patterns and species-specific regulatory programs can confound homologous cell-type alignment—remains to be systematically benchmarked, and may require incorporating evolutionary priors or species-aware regularization into the UOT formulation.

Several promising avenues remain for future exploration. Extending scPathOT to incorporate spatial transcriptomics (ST) and single-cell multi-omics (e.g., scATAC-seq) could allow the alignment of cellular states while preserving spatial proximity and epigenetic context. Moreover, the rapidly expanding landscape of large-scale perturbation atlases—including Perturb-seq[76], CRISPR-based genetic screens, and high-throughput drug perturbation datasets (e.g., sci-Plex[77], Tahoe-100M[78])— represents a natural application domain for scPathOT. Such datasets are typically characterized by heterogeneous perturbation-induced state shifts, substantial compositional imbalances across conditions, and the emergence of perturbation-specific cellular states, features that align closely with the design principles of our framework. Jointly modeling hundreds of perturbation conditions with scPathOT could facilitate the systematic identification of perturbation-responsive cellular states and support mechanistic hypothesis generation for therapeutic target discovery. Integrating our pathway-guided UOT mechanism with emerging single-cell foundation models[79] also holds potential to improve their adaptability to out-of-distribution disease datasets. As single-cell technologies continue to scale toward whole-organism atlases encompassing diverse perturbations, we anticipate that scPathOT, with its flexible, biologically informed integration capabilities, will contribute to the acceleration of translational biomedical research.

## Methods

### Data preprocessing and multi-view graph construction

#### Preprocessing and pathway activity estimation

Let the source domain (reference atlas) be 𝒟^s^ = **X**^s^, **y**^s^ and the target domain (query) be 𝒟^t^ = **X**^t^, where **X** ∈ ℝ^N×M^ is the raw count matrix of *N* cells and *M* genes and **y**^s^ contains source cell-type labels. Standard quality control was applied to remove low-quality cells and lowly expressed genes (thresholds listed in Supplementary Note 9). Counts were normalized to 10^6^ per cell (CPM) and transformed as ln(*x*+1).

From the preprocessed data we derive two complementary representations:

1. *HVG expression matrix*. The top *d*_hvg_ highly variable genes are selected by the dispersion method, yielding 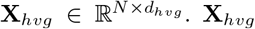 provides the node features shared across all graph views.
2. *Pathway activity matrices*. The framework accepts an arbitrary number of curated pathway databases as biological priors; in this study we use *V* = 2 databases, KEGG and Reactome, although additional resources such as GO, WikiPathways or MSigDB Hallmark can be incorporated without modifying the architecture. Each database *v* ∈ {1, …, *V*} contains *P*_v_ gene sets. For every database, AUCell is applied independently to score each of its pathways in each cell,

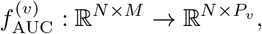

producing an activity matrix 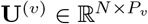 whose entry 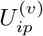 quantifies the enrichment of pathway *p* in cell *i*. The activity matrices are used only to define graph topology, never as node features.

#### Database-specific multi-view graph construction

Each pathway database is treated as a distinct biological view of the same cells. For view *v*, we build a cell– cell similarity graph whose topology is determined by activities in **U**^(v)^ rather than by raw expression. Treating each cell as a point in the *P*_v_-dimensional activity space of database *v*, we identify its *k* nearest neighbours by Euclidean distance (*k* = 5 throughout), giving a sparse binary adjacency

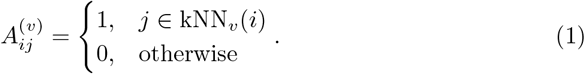

All *V* views share the same node feature matrix **X**_hvg_, so the data are represented as a set of graphs with shared features and view-specific topologies, 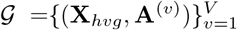. Because different databases emphasize different layers of biology— for instance, metabolism and signalling in KEGG versus reaction-level mechanisms in Reactome—each view captures cell–cell proximity along a distinct functional axis, and the downstream network learns to integrate them.

### Multi-view graph neural network encoder and dynamic fusion

### Graph neural network encoder with dual residual connections

Full-batch training on graphs of several hundred thousand cells is memory-prohibitive, so graph convolutions are executed on stochastic node-induced subgraphs. At each iteration we draw a mini-batch of *B* cells and retain only the edges of **A**^(v)^ whose endpoints both fall inside the batch, bounding the per-step edge cost by 𝒪 (*kB*) under *k*-regular kNN graphs.

For each view *v*, the shared node feature matrix 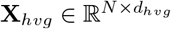 is first projected into a *D*-dimensional latent space,

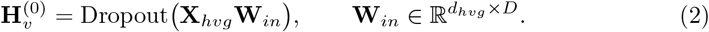

For *l* = 1, …, *L*, the representation is updated by

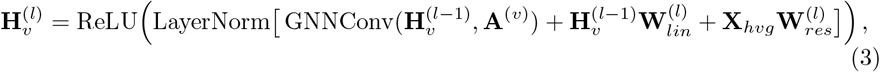

where GNNConv is a graph convolution operator (GCN, GAT or GraphSAGE, chosen per dataset; see Supplementary Note 9). Equation (3) combines two complementary residual paths: a layer-wise residual 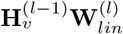 that follows the modern classic-GNN design of Luo *et al*.[80], and an initial residual 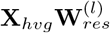 that re-injects the raw features into every layer to mitigate over-smoothing, following the GCNII scheme of

Rather than using only the last layer, we aggregate all *L* intermediate representations through a learned layer-wise gate. Concatenating 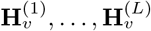 along the channel dimension gives 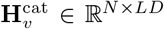, which is passed through a two-layer MLP with sigmoid output to produce feature-wise gates:

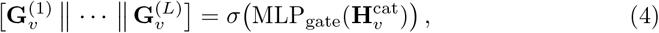

and the view-specific embedding is the gated sum

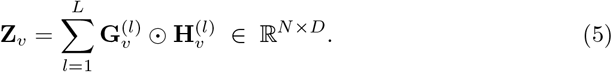

### Reconstruction auxiliary task

To regularize the latent space and discourage information loss in the encoder, we attach a view-specific decoder *D*_v_ (two-layer MLP with GELU activation and dropout) to every **Z**_v_ and reconstruct the original HVG profile,

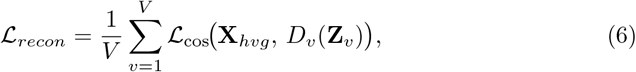

where ℒ_cos_ denotes the cosine embedding loss. Using cosine distance rather than mean-squared error emphasizes the relative co-expression structure of expression profiles and is less sensitive to scale differences across genes.

### Dynamic multi-view feature fusion

Different pathway databases contribute unequally to resolving different cell types, so the *V* view-specific embeddings are merged by a two-branch dynamic fusion module. Each **Z**_v_ is first passed through a shared feed-forward transform, following the standard Transformer FFN design,

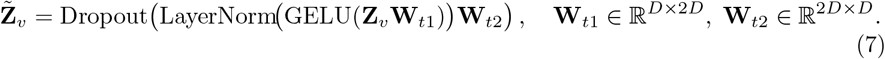

*(i) Cross-view self-attention*. Stacking the *V* transformed embeddings into **Z**_stack_ ∈ ℝ^N×V ×D^, we apply multi-head self-attention along the view axis and mean-pool over views to capture cross-view dependencies:

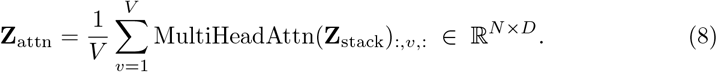

*(ii) Class-conditioned MLP gating*. In parallel, we compute per-cell weights over the *V* views. Let 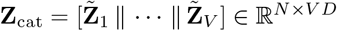 be the concatenation of transformed view features. A class-aware embedding 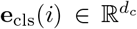 is obtained from a learnable lookup table 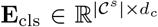 for source cells we use the ground-truth label, and for target cells—where no labels are available—we draw a uniform random index, which in practice acts as a stochastic regularizer that prevents the gate from overfitting to any particular class prior. The gating weights are then produced by a three-layer MLP with softmax output:

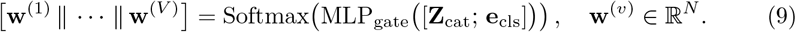

The gated feature is the weighted sum

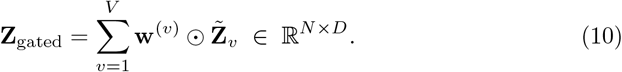

*(iii) Per-cell rescaling and bottleneck*. A per-cell attention score *α*_*i*_ = tanh(**z**_gated,*i*_ **W**_a1_) **W**_a2_ ∈ R (two-layer MLLP with tanh activation is combined with a scalar smoothing term 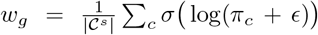 computed from the inverse-frequency class prior *π*_c_ of the source domain, yielding

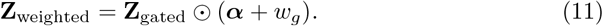

The rescaled feature is added residually to the cross-view branch and passed through a two-layer bottleneck MLP and *ℓ*_2_ normalization:

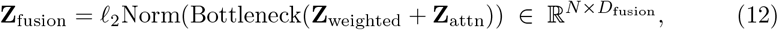

with *D*_fusion_ *< D* to yield a compact representation. The encoder and fusion module share parameters across source and target domains; we denote the resulting embeddings as **Z**^s^ and **Z**^t^, which serve as the input to the Stage 1 alignment objectives below. Unless otherwise stated we use *D* = 512, *D*_fusion_ = 256, *L* = 5 and *d*_c_ = 16; a full list of hyperparameters is given in Supplementary Note 9.

### Stage 1: Universal domain alignment and target-private (query-specific) class discovery

Given the fused representations 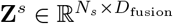 and 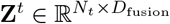, Stage 1 jointly fits a prototype classifier on the labeled source, (ii) aligns target cells of common classes to source prototypes through unbalanced optimal transport (UOT), and (iii) discovers target-private (query-specific) cell types by balanced OT clustering. The alignment and discovery objectives follow the UniOT paradigm of Chang *et al*.[8], adapted to the single-cell setting with a pathway-informed encoder and a scRNA-tailored adaptive filling strategy.

#### Source supervised classification

We parameterize a prototype classifier by a matrix of learnable source prototypes 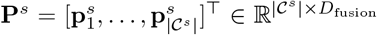, where 𝒞^s^ denotes the set of source cell types. Both cell features and prototypes are *ℓ*_2_-normalized at every forward pass; the prediction for source cell *i* is

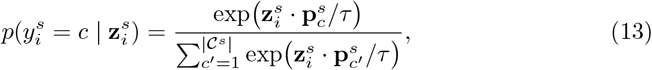

with temperature *τ* = 0.1. The supervised loss is the cross-entropy

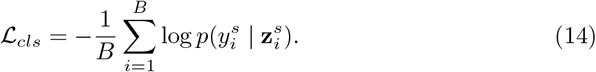

#### Target memory queue

OT solutions are sensitive to the marginal statistics of the input, which a single mini-batch estimates poorly. We therefore maintain a first-in-first-out memory queue 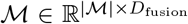 of size | ℳ | = 4*B* that stores the most recent normalized target embeddings. The queue is updated with the current batch at the end of every training step and serves two purposes: it provides a pool of background samples for the augmented UOT input below, and it is used to retrieve cross-batch nearest neighbours for the private class discovery module.

#### Common class detection via unbalanced OT

To align target cells from classes shared with the source while tolerating the presence of target-private cells, we cast cross-domain matching as an UOT problem. At each step, we draw 4*B* historical samples from ℳ and concatenate them with the current target batch to form an augmented feature matrix 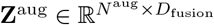 with *N* ^aug^ = 5*B*. The cosine similarity to source prototypes is

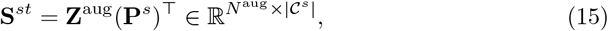

and we define the cost matrix **M**^st^ = − **S**^st^. Let 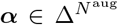 and 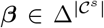 denote marginal priors over target samples and source classes, respectively. The UOT plan **Q**^st^ solves

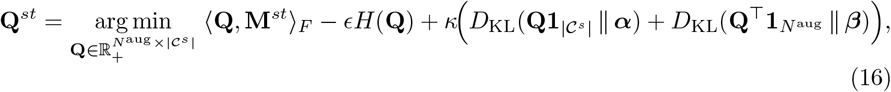

where *H*(**Q**) = − ∑ _ij_ *Q*_ij_ log *Q*_ij_ is the entropy regularizer[82] and *κ* controls the KL relaxation of the marginal constraints. In contrast to standard OT, the relaxed marginals allow mass destruction and creation, so target cells without a homologous source class need not be mapped onto any prototype. We solve Eq. (16) with the unbalanced Sinkhorn–Knopp algorithm implemented in the POT library[83], using *ϵ* = 0.03 and *κ* = 0.5.

The target prior is set to the uniform distribution 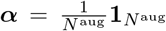. The source prior ***β*** is updated by exponential moving average across iterations:

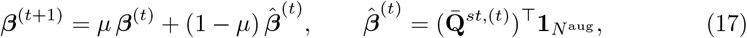

where 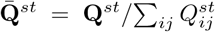 is the globally normalized plan, *µ* = 0.7, and 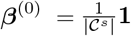. This adaptive prior prevents highly populated source classes from absorbing disproportionate mass.

#### Adaptive filling for batch imbalance

UOT is sensitive to the pseudo-positive/negative ratio within the input batch. We define the per-sample maximum similarity 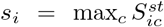 and classify sample *i* as a pseudo-positive if *s*_i_ *> γ* (we use *γ* = 0.7), otherwise as a pseudo-negative. Let *ρ*_pos_ denote the fraction of pseudo-positives in the augmented batch. Two regimes are handled differently:

i. *Positive filling (ρ*_pos_ ≤ 0.5). When pseudo-positives are under-represented, we synthesize additional positive rows from confident historical samples. We first run an auxiliary UOT pass over all *N* ^aug^ rows of **S**^st^ to obtain preliminary pseudo-labels and confident-sample indices. Inverse-frequency class weights **u** Δ are computed from the class distribution of these confident samples, and per-sample sampling weights are then obtained by indexing **u** with each sample’s pseudo-label. We multinomially resample *n*_fake_ = |𝒩_neg_ | − |𝒩_pos_ | rows (with replacement) from the confident set and prepend them to **S**^st^, producing the augmented similarity matrix used in Eq. (16).
ii. *Negative filling (ρ*_pos_ *>* 0.5). When pseudo-negatives are scarce, we fabricate virtual out-of-support samples. For every row *i* of **Z**^aug^, we identify its farthest source prototype 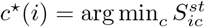 and construct a mixed feature

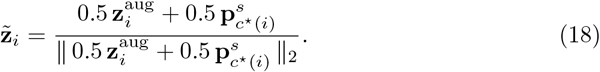

We uniformly sample *n*_fake_ =|𝒩_neg_ | − |𝒩_pos_ | of these mixed features and prepend their cosine similarities to **S**^st^ before solving Eq. (16).

#### Confidence-based pseudo-labeling

Let 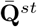 be the normalized UOT plan computed on the filled similarity matrix, and let 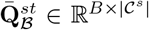 be its restriction to the rows corresponding to the current target batch (i.e., rows after the *n*_fake_ filled samples). For each target cell *i* in the batch we extract a confidence score and a candidate pseudo-label,

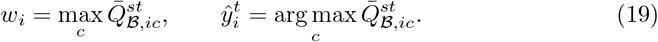

Because the total mass of 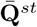 is one, a row whose mass is neither destroyed by UOT relaxation nor spread uniformly over classes satisfies *w*_i_ *>* 1*/N* ^aug^. We use this mass-conservation criterion as the confidence threshold; the selected index set is

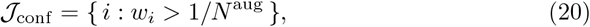

and the common class detection loss is a cross-entropy against the prototype classifier on these confident cells,

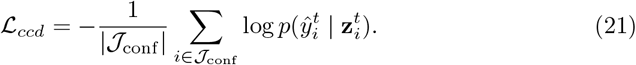

ℒ_*ccd*_ is activated only after a warmup period of 10% of the total training steps, during which the source classifier alone is optimized; this prevents pseudo-labels from propagating before the source prototypes stabilize.

#### Target-private class discovery

Target cells of novel classes are grouped by a second optimal transport problem with a separate set of learnable target cluster prototypes 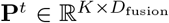, where *K* is set larger than |𝒞^s^| to accommodate unseen types. The cluster head produces *p*_cluster_(*k* | **z**) = Softmax(**z**(**P**^t^)^⊤^*/τ*)_k_.

At each step, we build a three-part augmented feature set combining (a) the current target batch (**Z**^t^, *anchor*), (b) for every anchor, a nearest neighbour retrieved from the current batch and ℳ(**Z**^nn^, *neighbour*), and (c) *B* background samples drawn uniformly from ℳ(**Z**^fill^). Concatenating gives 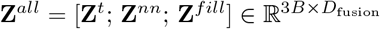, with similarity matrix **S**^tt^ = **Z**^all^(**P**^t^)^⊤^.

Unlike the UOT used for cross-domain alignment, clustering within the target domain benefits from strict marginal constraints to prevent degenerate solutions in which all samples collapse into a few clusters. We therefore solve the *balanced* OT problem

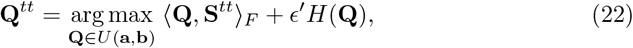

with 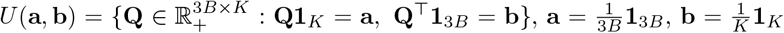, and *ϵ*^′^ = 0.05. The problem is solved by the Sinkhorn–Knopp algorithm[82].

Writing 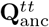 and 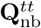 for the rows of **Q**^tt^ corresponding to the anchor and neighbour slices, and letting 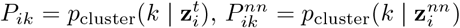, the private class discovery loss combines a cluster-assignment term and a swapped-prediction term in the spirit of SwAV[84]:

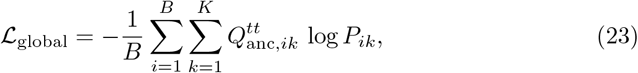

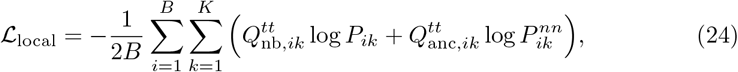

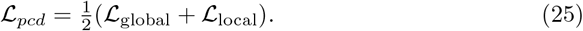

The swapped-prediction term enforces that an anchor and its nearest neighbour in ℳ agree on their cluster assignment, propagating local manifold information into the cluster prototypes.

#### Stage 1 objective

The four objectives are optimized jointly after the warmup period:

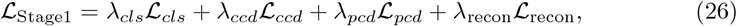

with default weights *λ*_cls_ = 1, *λ*_ccd_ = *λ*_pcd_ = *λ*_recon_ = 0.1. Source and target domains share all parameters up to the prototype layers, so the fused embeddings **Z**^s^ and **Z**^t^ produced at the end of Stage 1 reside in a single latent space that is (i) discriminative of known source classes, (ii) consistent across domains for common classes, and (iii) clustered for target-private (query-specific) classes. These embeddings, together with the UOT plan 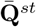, are the inputs to the Stage 2 knowledge distillation described next.

### Stage 2: Knowledge distillation for hard sample recovery

The UOT module of Stage 1 is deliberately conservative: the KL relaxation of marginal constraints destroys mass for any target cell whose embedding does not closely match a source prototype, and the confidence cutoff *w*_i_ *>* 1*/N* ^aug^ rejects borderline cells. As a result, cells with severe dropout, transitional phenotypes, or weak expression of lineage markers are frequently misclassified as target-private even though they belong to a common class. Stage 2 introduces a lightweight student network that recovers these misclassified cells by distilling the UOT plan while preserving genuine target-private (query-specific) calls.

#### Global UOT plan for distillation

After Stage 1 training terminates, we run a single global UOT pass over the entire target set using the trained encoder and frozen prototypes **P**^s^. In this final pass the memory queue is disabled (|ℳ| = 0 at evaluation), so the augmented target features consist only of the *N*_t_ real cells and *n*_fake_ filled samples introduced by the adaptive filling rule of Eq. 16, giving *N* ^aug^ = *N*_t_ + *n*_fake_. Let 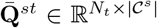 denote the globally normalized plan restricted to the real target rows; we use it for all subsequent Stage 2 computations.

#### Transport Outlier Score (TOS)

At each target cell *i*, the sum 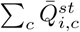 is the total mass that UOT assigns to the shared label set. Under a uniform target prior *α*_i_ = 1*/N* ^aug^, a cell whose embedding lies firmly in the source support retains this prior mass, whereas a cell whose embedding lies far from every source prototype has its mass relaxed away by the KL penalty. We define the Transport Outlier Score

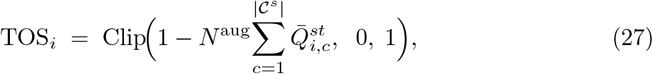

which measures the fraction of prior mass destroyed by UOT. TOS_i_ → 1 indicates lack of homology with any source class and therefore a high probability of being a genuine target-private cell or technical artifact (e.g., doublets); TOS_i_ → 0 indicates that the cell is well-matched to the source manifold. The Clip(·, 0, 1) is necessary because a small number of samples may absorb mass above their prior share when UOT concentrates transport.

**Teacher predictions and soft labels**. From 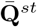 we extract three per-cell quantities used throughout Stage 2:

- the confidence score *w*_i_ = max_c_ 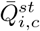;
- the open-set teacher prediction 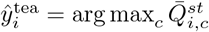 when *w*_i_ *>* 1*/N* ^aug^, else 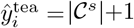 (marking the sample as target-private);
- the forced closed-set pseudo-label 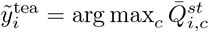, used only for class-balanced sample partitioning below.

Row-normalizing the source portion of 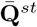 yields a soft label for each target cell,

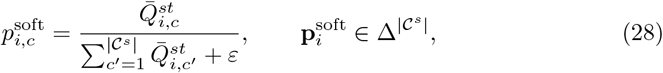

which serves as the distillation target for the student.

#### Tri-partition of target cells

We split 𝒟^t^ into three disjoint subsets that play distinct roles in distillation. Let *τ*_p_ = 0.7 be the outlier threshold and *γ*_anc_ = 0.6 the anchor quantile. For each source class *k* ∈ {1, …, | 𝒞^s^|}, let 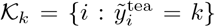 be the set of cells assigned to class *k* under the closed-set rule, and let *θ*_k_ be the top-*γ*_anc_ quantile of {*w*_i_ 𝒪:𝒪 *i* ∈ 𝒦_k_}.

*Outliers*: = {*i*: TOS_i_ *> τ*_p_ }. These cells have lost most of their prior mass under UOT relaxation and are treated as genuine target-private or technical outliers; they are frozen out of all distillation gradients.

*Confident set* 𝒜: Within each pseudo-class 𝒦_k_, a cell is admitted to 𝒜 if it satisfies all three conditions

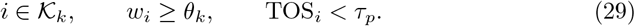

To protect minority cell types from being filtered out of every class, we override this rule for rare classes: whenever | 𝒦_k_| ≤ *δ*_rare_ (with *δ*_rare_ = 5), all cells in 𝒦_k_ are admitted to 𝒜 regardless of *θ*_k_ or TOS_i_. Using a class-specific quantile rather than a global threshold prevents long-tailed source distributions from starving rare classes of anchors. (The term “confident set” is used here to avoid collision with the PCD anchor–neighbour construction of Stage 1, which is unrelated.)

*Boundary cells* 𝒰: 𝒰 = 𝒟^t^ (𝒜 ∪ 𝒪). These cells are plausibly in-support (TOS_i_ is low) but the teacher was not confident enough to pass the class-specific cutoff. They are the main targets of the recovery mechanism.

#### Student network and dual-branch distillation

The student *S*_θ_ is a three-layer MLP (hidden widths 256 and 128, with BatchNorm, ReLU, and dropout 0.2) that maps the *frozen* Stage-1 embedding 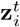 to a closed-set distribution 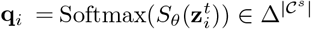. All encoder, fusion, and prototype parameters are held fixed during Stage 2; only *θ* is trained.

Two losses are applied to the two active subsets. On the confident set, the student mimics the teacher’s soft label through forward KL divergence,

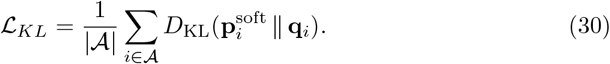

On the boundary set, where no reliable teacher signal is available, we apply entropy minimization,

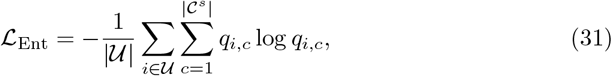

which sharpens the student’s predictions on cells lying in low-density regions of the decision boundary, a regularizer known to push boundaries away from high-density data regions and improve generalization under distribution shift[85]. Outliers in 𝒪 are excluded from both losses. The Stage-2 objective is

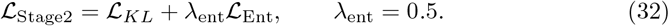

Training uses Adam with learning rate 10^−3^, weight decay 10^−4^, mini-batch size 256, for 200 epochs.

#### Inference-time recovery

At inference, the teacher and student are combined by a three-way decision rule. Writing 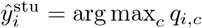, the final prediction is

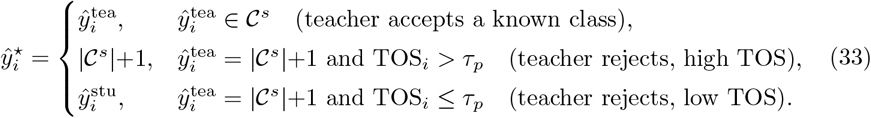

This rule assigns complementary roles to the two models: the teacher decides whether a cell belongs to the shared label set using the UOT-derived score TOS_i_, and the student predicts the specific class for cells that the teacher leaves at the margin. We do not apply an additional cutoff on the student’s softmax probability because the entropy-minimization term in ℒ_Stage2_ drives max_c_ *q*_i,c_ close to one for most target cells, rendering such a cutoff uninformative. The final output ŷ^⋆^ is the cell-type annotation reported by scPathOT.

### Downstream bioinformatics and statistical analysis

#### Annotation and integration metrics

Cross-domain annotation performance is evaluated by overall accuracy (ACC) and macro-averaged F1 score; the macro F1 is reported alongside ACC because it weights rare cell types equally with common ones, which matters for single-cell datasets with long-tailed class distributions. For atlas-level integration experiments, we report biological conservation (NMI, ARI, cell-type ASW, cLISI, isolated label F1, isolated label silhouette) and batch correction (batch ASW, graph iLISI, kBET, PCR batch, graph connectivity), all computed through the scIB benchmarking framework[5]. The overall integration score was computed as a weighted average of the biological conservation score and the batch correction score, with weights of 0.6 and 0.4, respectively, following the default scIB aggregation scheme.

#### Statistical analysis and enrichment toolkit

All statistical tests and visualizations were performed in Python. Differential expression was tested by Wilcoxon rank-sum through scanpy.tl.rank genes groups[86], with *p*-values adjusted by the Benjamini–Hochberg procedure (FDR *<* 0.05). Pathway enrichment was performed against the Enrichr gene-set libraries[87] using the gseapy Python client[88]: KEGG 2019 Mouse and Reactome 2024 for mouse data, and KEGG 2021 Human for human data. Cell–cell communication was analyzed with a Python re-implementation of the CellChat scoring scheme[89]: per-cell-type ensemble expression was summarized by the trimean, complex-subunit expression was aggregated by geometric mean, and directed ligand–receptor interaction strength between a sender type *i* and receiver type *j* was scored by the mass-action product *L*_i_ *R*_j_ on the human CellChatDB ligand–receptor database. Details of the implementation are provided in Supplementary Note 8.

#### Cross-disease cohort: functional heterogeneity of diabetic *β*-cells

Following annotation transfer from healthy to type 1 and type 2 diabetic donors, we analyzed functional heterogeneity of *β*-cells in the pathway-activity space learned by scPathOT. Pathways differentially active between healthy and disease *β*-cells were identified to reveal diabetes-specific dysregulation, and *β*-cells were clustered on the activity profile of these differential pathways to expose disease-associated functional subpopulations. Partition-based graph abstraction[90] (PAGA) was applied to the subpopulations to construct a topology-preserving state graph, along which pathway activity changes were tracked to characterize the molecular progression from the healthy baseline toward diabetic states.

#### Cross-age cohort: senescence scoring and co-expression networks

Following annotation transfer across mouse ages, senescence features of the bone marrow microenvironment were characterized per cell type using a *gene age-dynamic* (GAD) scoring scheme adapted from SenePy[53]. Age baselines were set adaptively: the young baseline was 3 months when the cell type contained more than 200 cells at that age and 1 month otherwise; the old baseline was 30 months when available and 24 months otherwise. Dynamic genes were retained at FDR *<* 0.05 and expression ratio *>* 1.5 over 1,000 permutations, and a co-expression network constructed by thresholding pairwise Pearson correlations was partitioned by Louvain clustering[91] to identify core gene hubs, defined as clusters whose within-hub connectivity significantly exceeded a random background. These hubs are reported as cell-type–specific senescence signatures; the GAD formula and full pipeline are given in Supplementary Note 7.

#### Cross-treatment cohort: immune microenvironment remodeling under chemotherapy

For the clinical pancreatic ductal adenocarcinoma cohort, annotations were transferred from the surgery-only (Non-NAC) cohort to the neoadjuvant chemotherapy (NAC) cohort to examine chemotherapy-induced remodeling of the tumor immune microenvironment. Per-sample abundances of major cell types were quantified, and fold-changes of NAC relative to Non-NAC were reported. Within each cell type, differentially expressed genes between the two groups were identified by Wilcoxon rank-sum and subjected to pathway enrichment to characterize the up-or down-regulated programs. From the ligand–receptor pairs linked to these differentially expressed genes, changes in interaction strength between NAC and Non-NAC were computed to construct a differential communication network summarizing treatment-induced shifts in intercellular signaling. Implementation details are provided in Supplementary Note 8.

## Acknowledgements

This work is supported by the National Natural Science Foundation of China [62433016].

